# Layer-specific spatiotemporal dynamics of feedforward and feedback in human visual object perception

**DOI:** 10.1101/2025.05.13.653501

**Authors:** Tony Carricarte, Siying Xie, Johannes Singer, Robert Trampel, Laurentius Huber, Zejin Lu, Tim C. Kietzmann, Nikolaus Weiskopf, Radoslaw M. Cichy

## Abstract

Visual object perception is mediated by information flow between regions of the ventral visual stream along feedforward and feedback anatomical connections. However, feedforward and feedback signals during naturalistic vision are rapid and overlapping, complicating their identification and precise functional specification. Here we recorded human layer-specific fMRI responses to naturalistic object images in early visual cortex (EVC) and lateral occipital complex (LOC) to isolate feedforward and feedback information signals spatially by their cortical layer-specific termination pattern. We combined these layer-specific fMRI responses with electroencephalography (EEG) responses for the same images to segregate feedforward and feedback signals in both time and space. Feedforward signals emerge early in the middle layers of EVC and LOC, followed by feedback signals in the superficial layer of LOC. Comparing the identified dynamics in LOC to a visual deep neural network (DNN), revealed that early feedforward signals in LOC encode medium-to-high complexity features, whereas later feedback signals increase the representational format to high complexity features. Together this specifies the spatiotemporal dynamics and functional role of feedforward and feedback information flow mediating visual object perception.

## Introduction

Human object vision relies on anatomical bidirectional connections along the ventral visual stream^1,2^, spanning the visual hierarchy from early visual cortex (EVC) to the lateral occipital complex (LOC)^3^. These connections mediate visual computations via feedforward and feedback information flows, with complex overlapping spatiotemporal dynamics^4–6^. While the feedforward flow carries sensory information up the visual processing hierarchy^7,8^, downstream feedback concurrently carries information down the hierarchy, refining and shaping feedforward neural dynamics^9^. Identifying the distinct contributions of these information flows to visual computation in space and time is therefore crucial for understanding the mechanism of human object vision.

Previous efforts to disentangle the specific signatures of feedforward from feedback have typically used one of two main approaches. The first involves invasive manipulations on the anatomically-^8^ and functionally-defined^6,10,11^ neural circuits in non-human primates, selectively targeting bottom-up vs top-down processes. While these interventions offer precise circuit-level control, their invasive nature makes them largely impractical for human research. The second approach comprises contrasting experimental conditions where the feedforward and feedback contributions vary, such as perception vs mental imagery^12^, attention vs non-attention^13^, early vs late backward masking^5^, and expectation vs sensory input^14,15^. However, this method does not directly assess information flow during naturalistic vision and has conceptual limitations, as it relies on indirect comparisons of experimental conditions that may be confounded by incompletely controlled differences between them^16–18^.

Instead, here we capitalized on the layer-specific anatomical connectivity found in the primate visual cortex^7,8,19^, to dissect feedforward from feedback signals: while feedforward connections terminate primarily in the middle layer^20^, feedback connections target superficial and deep layers^21,22^.

Based on this three-compartment model of cortical depth, we characterized the layer-specific spatiotemporal dynamics of feedforward and feedback signals in object perception. Using sub-millimeter resolution fMRI at 7T, we recorded layer-specific brain activity in human EVC, the initial stage of visual processing^23^, and in LOC, a high-level key region for object representations^3,24^, while participants viewed naturalistic object images. We then combined these layer-specific fMRI responses with time-resolved EEG responses for the same images within the framework of representational similarity analysis (RSA)^25–27^ to resolve feedforward and feedback processing in millisecond resolution. Based on this, we then characterized the representational format in terms of visual feature complexity of feedforward and feedback processing using artificial deep neural networks (DNNs).

## Results

Using sub-millimeter resolution 7T MRI, we recorded gradient-echo blood oxygenation level-dependent (GE-BOLD) signals in EVC and LOC of 10 participants in response to 24 different naturalistic object images (**Fig. 1A**). During the acquisition participants viewed naturalistic object images on real-world backgrounds in random order (**Fig 1B**). We estimated the neural response to each condition, i.e., object image, by fitting a general linear model.

**Figure 1.**
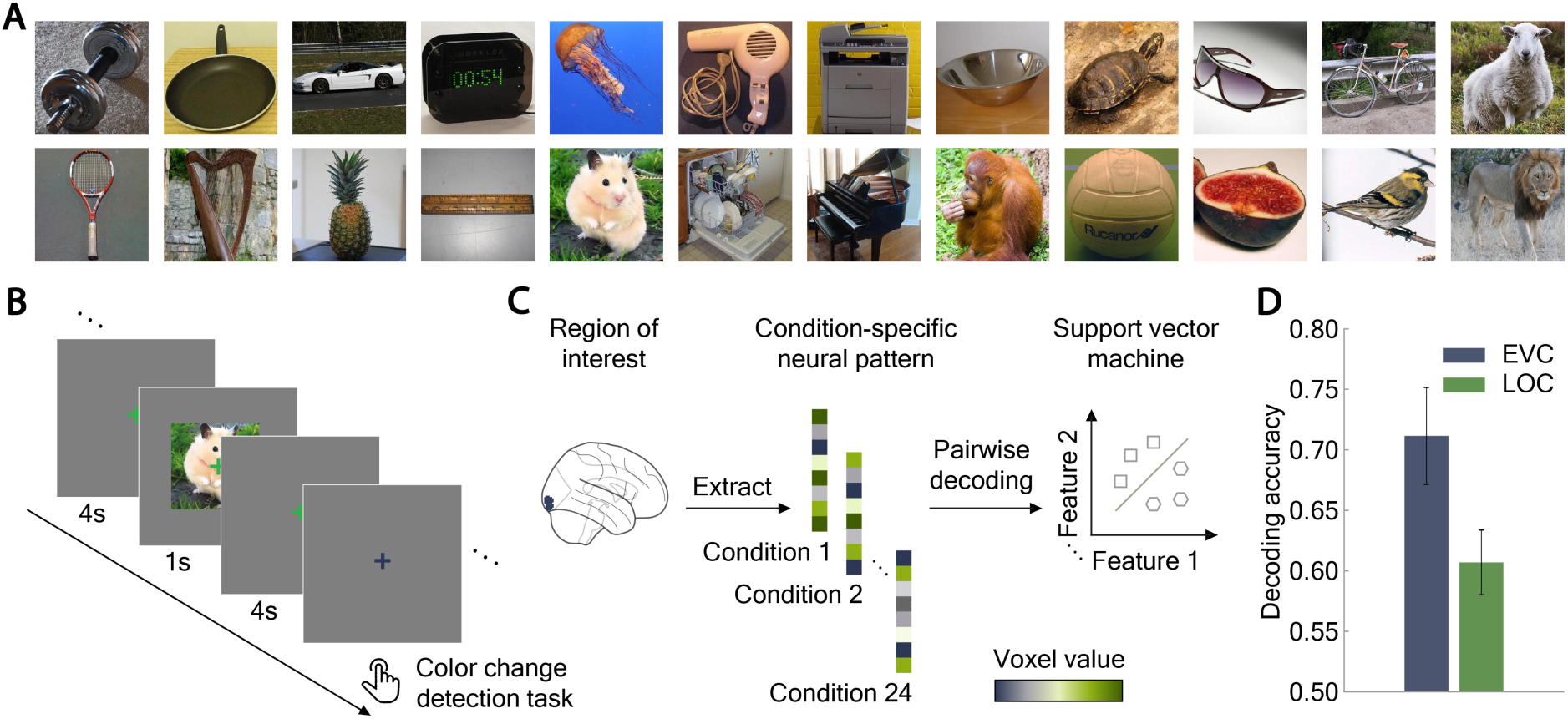
Stimuli, experimental design and multivariate pattern classification. **(A)** Stimulus set. The stimuli consisted of 24 different naturalistic object images. **(B)** fMRI experimental design. On each trial, participants viewed images for 1 s followed by a 4-s baseline interval. Participants were required to perform a color-change detection task on the fixation cross that occurred randomly throughout the experiment. **(C)** Extraction of voxel values to form condition-specific pattern vectors from region of interest (example here: EVC) and pairwise-object classification using a support vector machine. ROIs are depicted for visualization purposes only **(D)** Object-pairwise multivariate decoding output. Robust object-specific information was reliably decoded from EVC (71,16%, *P* = 0.0039) and LOC (60,69%, *P* = 0.0092) using one-sample permutation tests. Error bars indicate the standard error of the mean across participants.

Robust object information in both EVC and LOC is a precondition for further dissecting feedback and feedforward-related aspects. To ascertain this, we extracted voxel values from EVC and LOC separately to form condition-specific pattern vectors (**Fig. 1C**). Based on these pattern vectors we trained and tested a support vector machine (SVM) to perform pairwise-object classification of objects. We found robust decoding accuracy in both EVC (71,16%, *P =* 0.0039, one-sample permutation test) and LOC (60,69%, *P =* 0.0092), ascertaining reliable object information in those regions (**Fig. 1D**).

We then proceeded to identify and examine the spatiotemporal neural dynamics of visual representations in two steps that build upon each other. First, we assessed the macroscale of cortical regions (i.e., EVC and LOC) to establish and thus validate representational EEG-fMRI fusion at 7T, and to characterize the representational format of the identified dynamics. Based on this validation, we dissect feedforward from feedback neural processing at the finer mesoscale level of cortical layers.

### Spatiotemporal neural dynamics of object representations in EVC and LOC at the macroscale of brain regions

To establish the time course of object processing in EVC and LOC at the macroscale, we employed representational EEG-fMRI fusion^26–28^, which integrates time-resolved EEG with spatially resolved fMRI measurements for a combined spatiotemporally resolved view. For MRI, we used the dataset collected in this study, complemented with EEG responses from an existing dataset to the same set of 24 images^5^ as in the MRI experiment. We assessed the representational geometry of EVC and LOC with region-specific representational dissimilarity matrices (RDMs; **Fig. 2A**), and the representational geometry of EEG signals with RDMs in a time-resolved way from −200 to 600 ms with respect to image onset (**Fig. 2B**). We related EVC and LOC RDMs to the time-resolved EEG-RDMs, yielding a time course of representational similarity for EVC and LOC each, for which we report peak latency and peak latency differences with 95% confidence intervals in parenthesis as assessed by bootstrapping (1,000 iterations).

**Figure 2.**
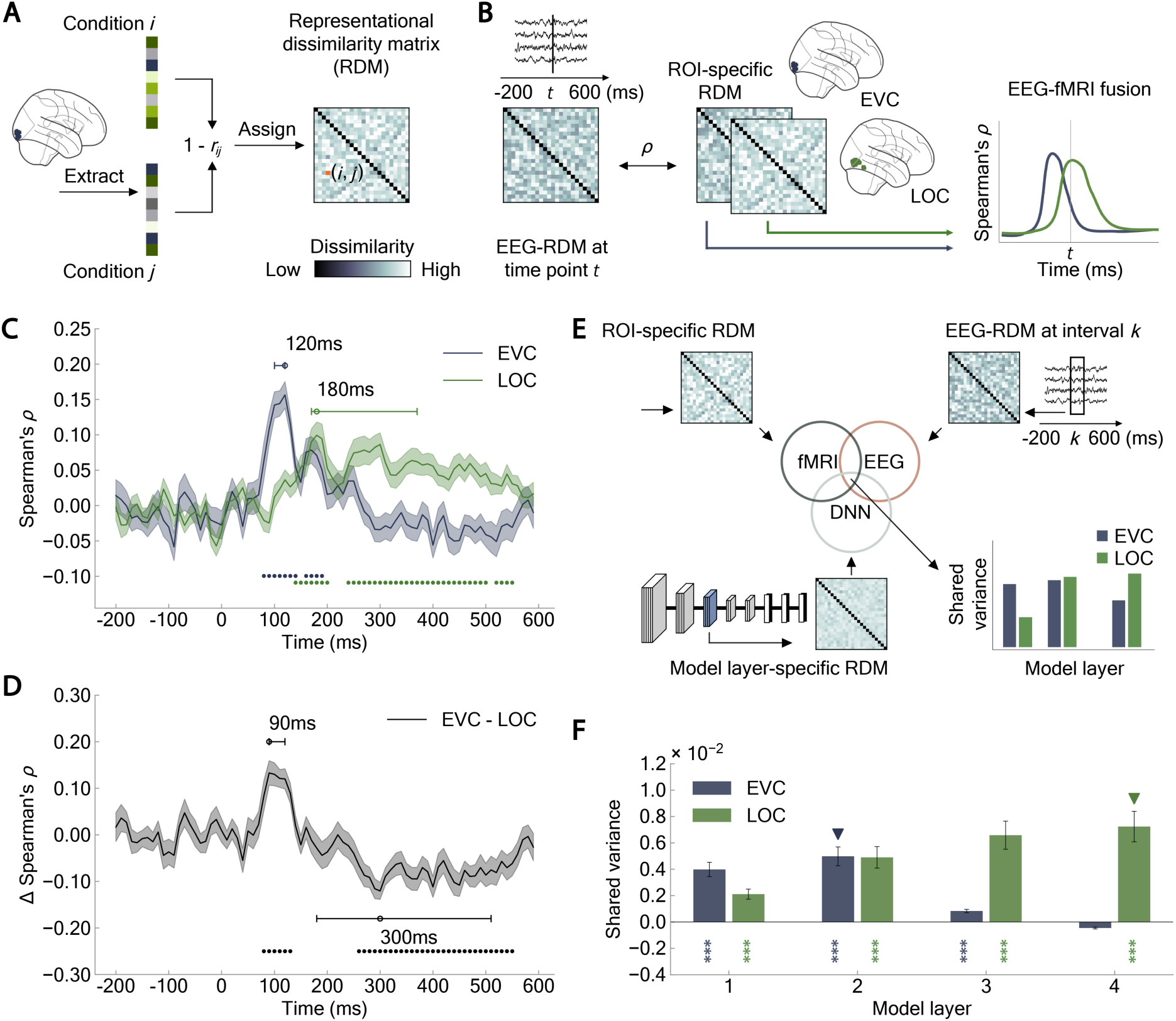
Macroscale representational EEG-fMRI fusion. **(A)** Representational similarity analysis. For each condition, we extracted the neural pattern from the region of interest (example here: EVC). We assessed the extent of pattern dissimilarity by calculating 1 - Pearson’s correlation for all combinations of experimental conditions (*i*, *j*) and assigned the dissimilarity values to an fMRI representational dissimilarity matrix (RDM) indexed by the conditions in rows and columns, at entry (*i*, *j*). ROIs are depicted for visualization purposes only **(B)** Representational EEG-fMRI fusion. For each time point *t*, we correlated the EEG-RDM to the fMRI-RDMs of EVC and LOC using Spearman’s rank order correlation. **(C)** Spatiotemporal neural dynamics at the macroscale level. Time course in EVC peaked earlier than in LOC. **(D)** Difference between EVC and LOC curves in **C**. EEG signals correlated first more with EVC than with LOC and later more with LOC than with EVC. **(E)** Commonality analysis. For each DNN layer, we correlated its RDM to each ROI-specific fMRI-RDM and the mean EEG-RDM at the time interval with significant ROI-specific temporal dynamics. **(F)** Format of representation (≈ feature complexity) in EVC and LOC. Visual representations of low-to-mid-level complexity emerge early in EVC, while mid-to-high-level object representations emerge later in LOC. Shaded regions and error bars indicate the standard error of the mean across participants. Colored circles denote significant time points (*N* = 32; cluster-defining threshold *P* < 0.05; cluster threshold *P* < 0.05); uncolored circles and horizontal lines indicate peak latency means and 95% confidence intervals, respectively. Colored asterisks indicate significant correlations (right-tailed permutation tests, FDR-corrected; \**P* < 0.05; \*\**P* < 0.01; \*\*\**P* < 0.001). Colored triangles represent model layers with the highest occurrence proportion, determined through 1,000-iteration bootstraps.

As expected from the well-established hierarchical organization of the visual system^7,8,29^, object processing in EVC preceded that in LOC (**Fig. 2C**): The EVC time course peaked at 120 ms (100 – 120 ms), whereas the LOC time course peaked later at 180 ms (170 – 370 ms), with a significant difference between peak times of 60 ms (50 – 250 ms; *P <* 0.001). Furthermore, in direct comparison by subtraction of the correlation curves of LOC from EVC, early representations correlated stronger with EVC than with LOC at 90 ms (90 – 120 ms; **Fig. 2D**), whereas late representations correlated stronger with LOC than with EVC at 300 ms (180 – 510 ms). A supplementary EEG-fMRI fusion analysis accounting for similarities in representational geometry between LOC and EVC by partial correlations confirmed these observations (**Suppl. Fig. 1A and B**), as also expected from the low correlation between the LOC and ECS RDMs (*R* = 0.05). Our results extend EEG-fMRI fusion^5,26,28,30^ from 3T to 7T fMRI and provide additional support to the procedure, thus warranting a spatially finer investigation of the spatiotemporal neural dynamics at the mesoscale level of cortical layers.

### The format of object representations in EVC and LOC at the macroscale of brain regions

The visual cortex represents objects in formats of increasing feature complexity from low to high along the ventral visual stream^31–34^. To confirm this progression here, we related the neural dynamics of EVC and LOC to the Vision Transformer (ViT) architecture^35^ from the DeiT family^36^ trained on object categorization^37^. The underlying rationale is that the feature complexity of visual representations progressively increases from lower to higher layers across the network’s layers, so that assessing the fit of the model layers to brain data reveals feature complexity of the neural representations^34,38^. We conducted commonality analysis, linking each of the 4 layer-specific DNN-RDMs to the ROI-specific fMRI-RDMs from EVC and LOC, and to the mean EEG-RDM at corresponding time intervals, defined by the significant time points of the raw curves in Fig. 2C. We report peak layers with 95% confidence intervals in parenthesis (1,000 bootstraps).

We found commonality with predominantly low to mid model layers, with a peak at model layer 2 (1 – 2) in EVC. and with predominantly middle to high model layers with a peak at model layer 4 (3 – 4) in LOC (**Fig. 2F**). This indicates representations of primarily low- to mid-complexity in EVC and mid- to high-complexity in LOC. This pattern of results remained consistent when using an alternative experimental choice based on significant time points from the difference between EVC and LOC curves in Fig. 2D (**Suppl. Fig. 1C**). Our results thus confirm a transition from representations of low-to-mid complexity processed early in EVC into representations of higher complexity processed later in LOC^28^, further providing additional support to the analytic approach at 7T and warranting a finer investigation of representational format at the mesoscale level.

### Spatiotemporal neural dynamics of object representations in EVC and LOC at the mesoscale of cortical layers

To investigate the spatiotemporal neural dynamics at the mesoscale level, we applied the research procedure validated above to the finer level of cortical layers (**Fig. 3A**). To this end, we segmented the cortical ribbon into three equidistant layers^39^: deep, middle and superficial. We then applied representational EEG-fMRI fusion as established above, but now at the level of layers, to yield time-resolved and layer-specific visual object processing time courses in EVC and LOC. To control for non-specific macrovascular responses^40^ that affect layer specificity^41^ and are reflected as increased GE-BOLD responses towards superficial layers (**Suppl. Fig. 1D**), we conducted EEG-fMRI fusion analysis partialing out for each layer the effect of the layers beneath.

**Figure 3.**
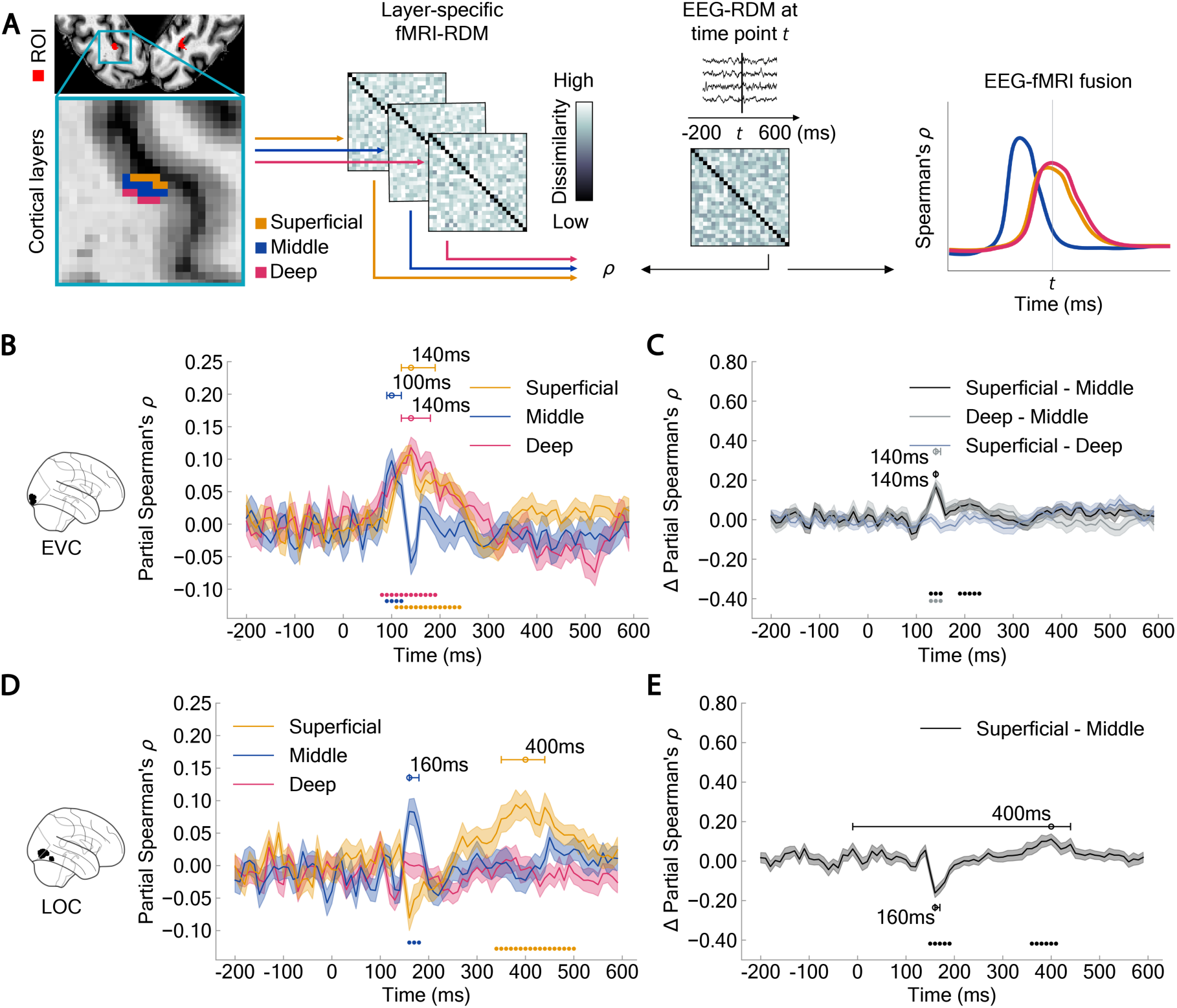
Mesoscale representational EEG-fMRI fusion. (**A**). For each cortical layer — deep, middle, and superficial — (example here: for EVC) we computed the partial Spearman’s rank-order correlation between its layer-specific fMRI-RDM and the EEG-RDM at each time point *t*. **(B)** Layer-specific spatiotemporal neural dynamics in EVC. EEG signals correlated early across layers, with the time course in the middle layer peaking earlier than in the deep and superficial layers. **(C)** Difference between EVC layer curves in **B**. EEG signals correlated lately more with deep and superficial layers than with the middle layer **(D)** Layer-specific spatiotemporal neural dynamics in LOC. EEG signals correlated early in the middle layer and later in the superficial layer. **(E)** Difference between LOC layer curves in **D**. EEG signals correlated early more with the middle layer than with the superficial layer, and later more with the superficial layer than with the middle layer. Shaded regions indicate the standard error of the mean across participants. Colored circles denote significant time points (*N* = 32, cluster-defining threshold *P <* 0.05, cluster threshold *P <* 0.05); uncolored circles and horizontal lines indicate peak latency means and 95% confidence intervals, respectively.

In EVC we observed a correlation pattern suggesting two processing stages with a common rise (**Fig. 3B, C**) indexed by different profiles across layers and in timing. The first stage is marked by a peak in the middle layer at 100 ms (90 – 120 ms; **Fig. 3B**). The second stage is characterized by peaks at 140 ms in both the deep (120 –180 ms) and superficial (120 – 190 ms) layers, with significant latency differences of 40 ms (10 – 50 ms; *P =* 0.004) between the middle layer and the deep layer and 40 ms (20 – 80 ms; *P =* 0.002) between the middle layer and superficial layer, but not between the deep layer and the superficial layer 0 ms (−20 – 40 ms; *P =* 0.814). This pattern was further substantiated by subtracting the layer-specific time courses which showed significant effects (**Fig. 3C**). We observed stronger correlations of late representations at 140 ms with the deep (140–150 ms) and the superficial (140–140 ms) layers than with the middle layer. This suggests initial feedforward processing in the middle layer, followed by later signals in the deep and superficial layers that could reflect either feedback processing or the sequential propagation of feedforward activity along the canonical microcircuit^8,42^ in EVC.

In LOC, the results pattern also indicated two stages with a distinctive layer and temporal profile (**Fig. 3D, E**). We observe earlier processing in the middle layer, followed by processing in the superficial layer later (**Fig. 3D**). In detail, time courses of object processing peaked first in the middle layer at 160 ms (160 – 180 ms), and later in the superficial layer at 400 ms (350 – 440 ms), with a significant difference in peak latency of 240 ms (110 – 290 ms; *P <* 0.012). A direct comparison of time courses by subtraction (**Fig. 3E**) substantiated the observation in that representations correlated stronger with the middle layer early than the superficial layer at 160 ms (160 – 170 ms), and late representations correlated stronger with the superficial layer than the middle layer later, at 400 ms (90 – 440 ms). This suggests an initial feedforward processing in the middle layer and a later emerging feedback processing with a distinctive layer profile in the superficial layer in LOC.

An analogous results pattern in EVC (**Suppl. Fig. 2A, C**) and LOC (**Suppl. Fig. 2B, D**) was observed when assessing layers directly without partialing out the effect of deeper layers. To ensure that differences in the number of voxels across cortical layers did not bias the representational dissimilarities, we repeated the analyses using an equal number of voxels per layer by randomly subsampling to match the layer with the lowest voxel number. This control analysis yielded comparable results in EVC (**Suppl. Fig. 2E**) and LOC (**Suppl. Fig. 2F**), confirming the robustness of the results to particular analysis choices.

Together, this resolves the spatiotemporal dynamics of feedforward and feedback information flow during visual object processing across EVC and LOC through cortical layer-specific and temporally distinct response profiles.

### The format of object representations in EVC and LOC at the mesoscale of cortical layers

One interpretation of the observed layer-specific spatiotemporal dynamics, being guided by the functional pattern of cortical layer connectivity^43,44^, is that they indicate interareal communication via feedforward and feedback connections^42,45^. However, an alternative explanation is that they arise intra-areal communication via lateral connections^46^ or from superficial bias in the fMRI measurements^47^.

To disambiguate between these options, we characterized the format of representations by assessing feature complexity as present in DNN layers, from low to high, applying the research approach as used at the macroscale but now at the level of cortical layers.

Feedforward processing within a cortical region involves convolution-like computations from simple to complex and hypercomplex cells^48,49^, progressively transforming representations to achieve some tolerance. We conceptualize this as a transformation that changes the tolerance of a feature, not the intrinsic feature complexity itself. Instead, an increase in intrinsic feature complexity requires the integration of different features into new, emergent, and more elaborated representations, as those reflected across different DNN layers. Feedback processing, in contrast, is likely to play an active role in this feature complexity increase^5,50^ by transferring higher-complexity representations. Accordingly, if the observed layer-specific spatiotemporal dynamics reflect an interplay of feedforward and feedback information flow, we would expect the flow to carry information of varying complexity^51,52^ across different levels of the visual hierarchy^53,54^, resulting in distinct representational formats across cortical layers.

We conducted commonality analysis, linking each model layer-specific DNN-RDM to the deep, middle and superficial layer-specific fMRI-RDM and mean EEG-RDM at corresponding time intervals, defined by the significant time points of the raw curves in Fig. 3B and D indicating periods of layer-specific neural dynamics (**Fig. 4A**).

**Figure 4.**
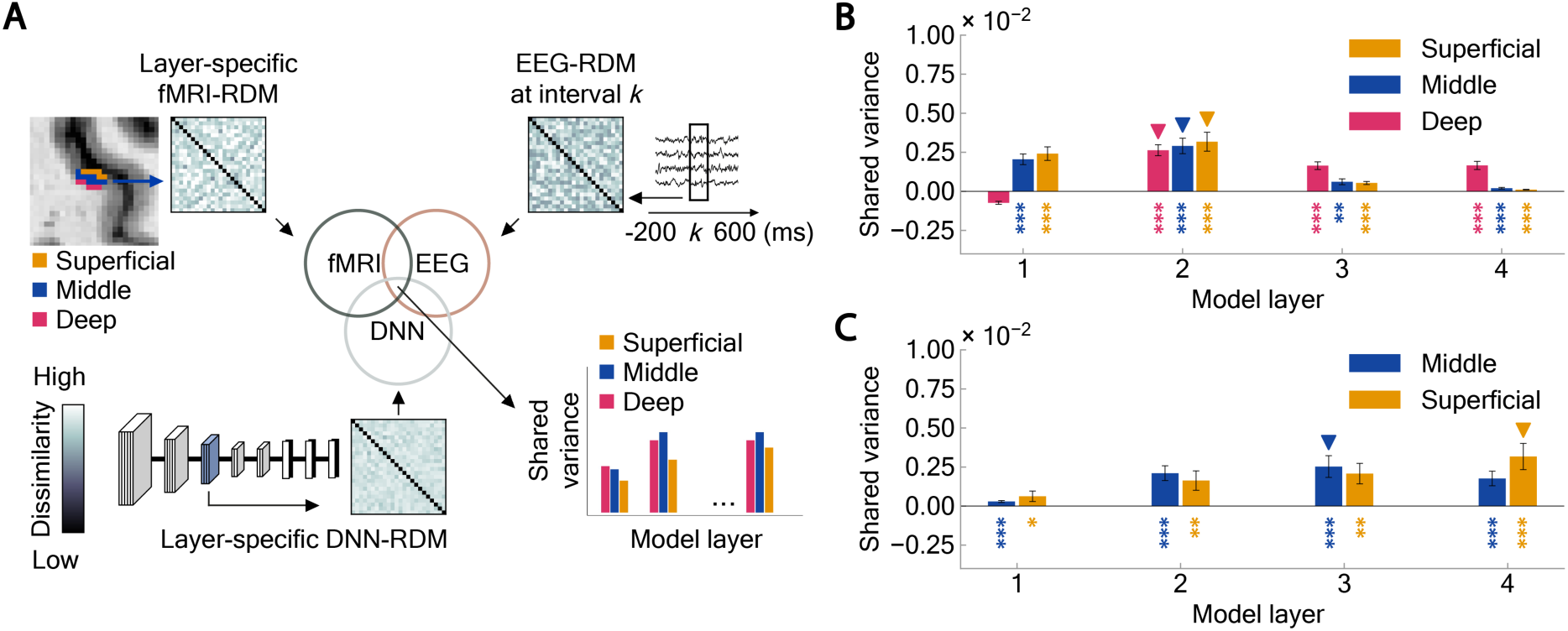
Mesoscale commonality analysis between fMRI, EEG and ViT. **(A)** Procedure. For each transformer layer, we correlated its RDM to each layer-specific fMRI-RDM in EVC and LOC and the mean EEG-RDM at the time interval with significant layer-specific temporal dynamics. **(B)** Format of representation (≈ feature complexity) across cortical layers in EVC. Low model layers correlated strongly across layers in EVC. **(C)** Format of representation (≈ feature complexity) across cortical layers in LOC. Middle model layers correlated strongly with the middle layer in LOC, while high model layers correlated primarily with the superficial layer. Colored asterisks indicate significant correlations (*N* = 32, right-tailed permutation tests, FDR-corrected; \**P <* 0.05; \*\**P <* 0.01; \*\*\**P <* 0.001). Colored triangles represent model layers with the highest occurrence proportion, determined through 1,000-iteration bootstraps.

In EVC (**Fig. 4B**) we observed a uniform results pattern across cortical layers: object representations shared variance with all DNN layers, but most strongly with model layer 2 (2 – 2), indicating mainly representations of low-to-mid level complexity. This suggests that the observed layer-specific spatiotemporal dynamics with a common rise in EVC reflect feedforward propagation, lateral connections modulating the neural gain^55–59^ or superficial bias^47^, rather than feedback.

In contrast, in LOC (**Fig. 4C**) we observed a shift from mid- to high feature complexity across cortical layers. The early emerging representations in the middle cortical layer (**Fig. 3D**) were of mid-to-high-complexity, as indicated by a peak at model layer 3 (2 – 3). In contrast, the late emerging representations in the superficial layers were of high complexity, with a peak at model layer 4 (4 – 4). A direct comparison of feature complexity between the middle and superficial layers, using bootstrapped peak layer differences, confirmed that the superficial layer representations encoded higher-level features than the middle layer (1 model layer difference; *P* < 0.001).

These results proved robust across multiple experimental choices. These included using an equal number of voxels per layer (**Suppl. Fig. 2G and H**), testing a different network architecture, AlexNet^37^ (**Suppl. Fig. 2I and J**) to account for DNN-specific biases, averaging over significant time intervals from the layer-curve differences from the layer-curve differences in Fig. 3C and E (**Suppl. Fig. 2K and L**), and analyses across individual time points spanning the full time course (**Suppl. Fig. 3**).

Together, these findings suggest a dynamic shift in representational format in LOC, transitioning from early representations of mid-level feature complexity in the feedforward flow to later representations of high feature complexity through interareal feedback.

## Discussion

We leveraged layer-resolved fMRI, time-resolved EEG data and DNNs to identify and characterize the spatiotemporal neural dynamics of feedforward and feedback information flow underlying object perception.

We validated our methods using 7T fMRI at the macroscale of cortical regions by replicating the temporal dynamics and representational format in EVC and LOC observed in 3T fMRI studies^5,26,30,34^, allowing us to proceed to the mesoscale of cortical layers. There we made two key observations. First, we observed distinct layer-specific temporal profiles for EVC and LOC. Visual representations in the middle layers emerged earlier than in the deep and superficial layers of EVC and the superficial layers of LOC. Second, the identified layer-specific dynamics in LOC had distinctive visual feature complexity profiles: the early emerging middle layer representations were of mid-to-high complexity and the later emerging superficial layer representations were of high complexity.

### Layer-specific EEG-fMRI fusion reveals sequential feedforward and feedback processing in visual object perception

Visual object perception unfolds through intricate spatiotemporal neural dynamics^4–6,60,61^, mediating feedforward and feedback information flow^26,62^ rapidly and through temporally overlapping responses. Here, we disentangle the temporal dynamics of feedforward from feedback signals by leveraging the anatomical canonical microcircuit of cortex^7,8^ at the input and output stage of cortical object processing^3,63,64^ – EVC and LOC. We find that feedforward signals emerge early in the middle layer of both EVC at 100 ms and in LOC at 160 ms, while feedback appear relatively later in superficial layer of LOC at 400 ms, supporting the idea of sequential processing of visual information first through feedforward, then through feedback processing^44,65^. Our results provide a functional temporal characterization for the anatomical canonical cortical microcircuit model of feedforward and feedback connectivity in the human ventral visual stream.

Although the observed differences in feedforward peak latency between EVC and LOC indicate a consistent temporal hierarchy, their interpretation warrants caution due to the indirect nature of the fusion-derived measures. Note that there is no trivial way to relate the exact peaks reported in the fusion analysis to other measures of temporal dynamics in cortex, such as spike timing^11,66^. They reflect the relation mass signals in the EEG with BOLD signals in the fMRI, involving the foveal confluence and thus aggregated across V1 to V3. Thus, while the timing and in particular its differences across regions and layers as determined by fusion are robust and define a clear order, the relationship to neural activity measured invasively in the brain needs further confirmation.

While previous studies have consistently reported feedback modulation in the EVC^11,54,67–69^, our results suggest that the observed layer-specific spatiotemporal dynamics instead reflect feedforward propagation, lateral interactions modulating the neural gain^55–59^ or superficial bias^47^, with no evidence of interareal feedback. This apparent lack of feedback signals could reflect the limited laminar specificity of our measurements, influenced by vascular contamination and partial volume effects^47,70^ (see limitation section for more details), effectively masking feedback signals under the prevailing feedforward activity in the EVC.

Our findings have theoretical implications. For example, they specify in spatiotemporal terms the dynamical communication model in predictive coding theory^71^ which posits distinct neural channels for the transmission of sensory and predictive information^72–74^. Our results indicate that whereas sensory signals are conveyed early in middle layers, subsequent predictive signals are transmitted later as feedback in superficial layers. This specification opens a new path for further testing predictions of predictive coding by investigating the content and integration of predictive feedback and sensory feedforward signals^75–77^. For example, our results invite investigating layer-specific neural dynamics at distinct frequency bands for feedforward and feedback processing as predicted from human^78,79^ and non-human invasive studies^72,80–82^, or recently the spatiotemporal dynamics of false percepts^73,74^.

Our approach facilitates resolving the interplay of feedforward and feedback processing in a variety of visual research contexts. It may allow dissecting the distinctive roles of feedforward and feedback information flow crucial to cognitive functions such as attention, expectation and memory by identifying and characterizing their distinct spatiotemporal dynamics. Similarly, using multi-rather than univariate methods^12,72,83^ to assess layer-specific fMRI extends content-sensitive analysis^84–86^ from the macro- to the mesoscale of cortical organization, allowing for the assessment of the representational contents of feedforward and feedback information flow underlying human cognition.

### Layer-specific EEG-fMRI fusion clarifies neural dynamics of high-level visual cortex regions

Neural dynamics in response to faces^26^, scenes^87^ and objects^88,89^ assessed at the macroscale of cortical regions commonly display a double peak pattern (see also **Fig. 2C**), with a sharp, early peak around 160-200 ms followed by a wide second peak around 240-450 ms, suggesting distinct contributing neural circuits and kinds of processing. Our layer-specific results clarify the neural circuits^26^ and functional nature of the components underlying this pattern: it results from mixing the early, narrow-peaked and neural dynamics at the middle layer ∼160 ms of feedforward information flow with the later, wide-peaked and thus more persistent dynamics at the superficial layer ∼400 ms of feedback information flow. Whereas the early peak latency matches that of sensory feedforward signals^90^, the late peak dynamics match that of and attention-^91^, consciousness-^92^ or task-related^93^ feedback, originating from higher-order brain regions, such as the frontal eye fields^10^ or the prefrontal cortex^94^, outside the visual cortex. Thus, our results provide circuit-level mechanistic as well as functional interpretative guidance for standard 3T human neuroimaging studies that typically cannot resolve visual information flow at this fine spatiotemporal level.

We did not observe sustained activity in EVC following the middle-layer peak observed in LOC. This difference may reflect variations in temporal processing across the visual hierarchy during object perception, potentially arising from distinct contributions of feedforward and feedback information flow. Specifically, neural dynamics in EVC are likely dominated by rapid, sensory-driven feedforward input, whereas LOC exhibits both feedforward and more sustained feedback components. Such prolonged activity has been reported previously in higher-level visual areas^26,88,89^, but not in EVC. We therefore do not interpret the absence of sustained EVC activity as a lack of feedback per se, but rather as a limitation of the spatial and temporal sensitivity of our measurements, which likely capture the dominant population-level dynamics, consistent with patterns observed in univariate analyses^95^.

### Differential representational format across layer-specific dynamics in LOC implies interareal feedback mediating high-complexity visual features

The analysis of the representational format of the neural dynamics in LOC suggests that early feedforward signals primarily convey mid-complexity features, while feedback signals convey high-complexity features. This is consistent with the idea that feedback does not merely modulate the gain of pre-existing features but is actively involved in the emergence of additional, more complex features that are absent in feedforward processing^5^. This finding supports the notion that interareal feedback is integral to core object recognition^96–98^. A potential source of this feedback might be the dorsolateral prefrontal cortex (DLPFC)^99^, whose silencing decreases feature complexity in monkey inferior temporal cortex^50^, the homologue of human LOC. Based on our results we predict that disrupting processing in human DLPFC, e.g., through transcranial magnetic stimulation^100,101^, will yield analogous effects specifically on superficial, late cortical layer responses in LOC.

### Limitations of the study and future research

We based our analyses on the GE-BOLD signal that is affected by locally nonspecific responses from macrovasculature^47^, compromising the estimation of the laminar response. To mitigate this effect while assessing the relationship between neural patterns, we applied a two-steps approach. First, we used Pearson correlation, which is invariant to signal scaling and thus less sensitive to pattern amplification arising from differences in vascular density across the cortical ribbon^102^. Second, we applied partial correlations to remove residual baseline effects originating from underlying layers. Although this approach enabled us to assess pattern relationships reducing unwanted blood contamination effects, non-linear signal leakage, particularly from draining veins toward the pial surface might still persist^103^.

To ensure sufficient signal to noise ratio (SNR) while covering both EVC and LOC, we used an isotropic voxel size of 0.9 mm. However, given that the cortical thickness in the visual cortex often does not exceed three times this resolution^104^, partial volume effects might exist, resulting in signal mixing across cortical layers.

Despite these limitations, our methodological framework holds exciting potential for noninvasive, in vivo resolution of the spatiotemporal dynamics of feedforward and feedback processing in the human brain. Here, we combined EEG and fMRI data to capture common and stable neural effects across modalities using a condition-by-condition approach, whereas other studies have employed trial-by-trial methods^105^. Exploring how these two approaches could be integrated offers an interesting avenue for future research, as it may provide complementary insights into the coupling between electrophysiological and hemodynamic signals.

Key challenges remain, including expanding spatial coverage without compromising signal quality, which would enable the study of the spatiotemporal dynamics of object information flow across the entire ventral visual stream and beyond. Future human fMRI studies using acquisitions with higher spatial specificity^106–108^, optimized for condition-rich experimental designs^25^, will be essential to confirm and extend our findings.

## Conclusion

Understanding how the abundant feedforward and feedback connections in visual cortex mediate object vision requires specifying their functional role. Here we provided two key advances in this regard. First, leveraging layer-specific fMRI with EEG-fMRI fusion we dissociated temporally overlapping feedforward and feedback processing in LOC and EVC, specifying their unique temporal profile. Second, by assessing feature complexity, we showed that feedback in LOC might actively increase the complexity of LOC’s feature format.

## Acknowledgments

This work was funded by the German Research Foundation (DFG, CI241/1-1, CI241/3-1, INST 272/297-1 to R.M.C.), by the European Research Council (ERC) starting grant (ERC-2018-STG 803370 to R.M.C.), supported by the NIH Intramural Program of NIMH/NINDS (#ZIC MH002884 to R. H.) and by the Einstein Center for Neuroscience (to T.C.). Computing resources were provided by the high-performance computing facilities at ZEDAT, Freie Universität Berlin, Germany.

## Materials and Methods

### fMRI participants

10 adult volunteers (mean age 29.4 years; age range 20-37 years; 5 female) participated in the study and provided written informed consent. The sample size was based on previous conventional layer fMRI studies conducted at 7T^83,106,109^. All participants had normal or corrected-to-normal vision and no history of neurological disorders. All participants received a monetary reward at the end of the study. The study was approved by the Ethics Committee of the Faculty of Medicine at Leipzig University, Germany, and conducted in accordance with the ethical principles of the Declaration of Helsinki, except for study preregistration, which was not performed.

### Stimulus set

The stimulus set consisted of 24 naturalistic color images of everyday objects on real-world backgrounds, each of a different category from the ImageNet image database (i.e., animals, tools, vehicles, foods and others)^34^.

### fMRI experimental design

The study was composed of a main experimental part and one localizer run.

The main experimental part consisted of 6 to 14 runs (mean ± SD: 9.9 ± 3.2), each lasting 621 s. Each run started and ended with a 10.5-s baseline period and included all 24 images, each repeated five times, for a total of 120 trials presented in random order. Each trial began with a 1-s stimulus-on interval, during which a centrally displayed object image (5° visual angle) appeared on a grey background with a pink fixation cross. We used a short stimulus-off interval of 4 s and randomization of condition ordering, consistent with previous RSA studies^5,25,26,28^, to allow for both a high number of conditions and trial repetitions with the goal of high reliability of the estimated activity patterns (**Suppl. Fig. 4A, B and C**). During this interval, only the fixation cross was displayed. Participants were instructed to maintain their gaze on the fixation cross throughout the experiment and to perform a color-change detection task unrelated to the object images, thereby, minimizing task-dependent effects during object processing^110^. Participants responded by pressing a button as soon as the fixation cross turned blue for 300 ms.

The functional localizer run was intended to define regions of interest (ROIs) EVC and LOC. Participants completed it at the beginning of the recording session. The localizer consisted of 15-s blocks displaying objects (not included in the main experiment) and scrambled objects overlayed with a fixation cross, interleaved with 7.5-s baseline blocks showing only a fixation cross on a grey background. In each block images were centrally presented (12° visual angle) for 400 ms, followed by a 350-ms display of the fixation cross. Participants were instructed to maintain their gaze on the fixation cross and to press a button if the same image appeared in consecutive trials. The localizer run included 12 blocks of each image type, resulting in a total duration of 465 s. Block order was pseudo-randomized to avoid immediate repetition of the same block type.

### MRI Procedure

MRI data were acquired at the Max Planck Institute for Human Cognitive and Brain Sciences in Leipzig, Germany. Four participants completed the experiment in one scanning session. Six participants completed the experiment in two scanning sessions on two separate days. During the first scanning session, we acquired a T1-weighted anatomical image, the functional localizer and 5 to 8 runs of the main experiment. During the second scanning session, participants completed 6 to 8 runs of the main experiment. Additionally, to enable distortion correction, five volumes with reversed phase-encoding polarity were acquired following the first run of each main experimental session. To ensure that participants were familiar with the experimental tasks, we provided them with verbal and written instructions prior to the scanning, and the participants completed a 2-min training for both the localizer and the main tasks.

### MRI acquisition parameters

We acquired MR images on a Siemens Magnetom Terra 7T whole-body system (Siemens Healthineers, Erlangen, Germany) with a single-channel-transmit and a 32-channel radio-frequency (RF) receive head coil (Nova Medical Inc, Wilmington, USA). We acquired the functional data using a 2D Gradient-echo (GE) echo planar imaging (EPI) sequence^111^ (voxel size = 0.9 mm isotropic resolution, TE/TR = 26.2/3500 ms, in-plane field of view (FoV) 192 × 192 mm^2^, 48 axial slices, flip angle = 75°, echo spacing = 1.0 ms, GRAPPA factor = 3, partial Fourier = 6/8, phase encoding direction anterior-posterior). We recorded anatomical data using an MP2RAGE sequence^112^ (voxel size = 0.7 mm isotropic resolution, TE/TR = 2.01/5590 ms, in-plane FoV 224 × 224 mm, GRAPPA factor = 2) yielding two inversion contrasts (TI1 = 900 ms, flip angle 1 = 5°; TI2 = 2750 ms, flip angle 2 = 3°). The two inversion contrasts were combined to produce T1-weighted MP2RAGE uniform (UNI) images with high contrast to noise ratio.

### MRI preprocessing

For each recording session, we spatially realigned the functional volumes to their mean volume using SPM12 (http://www.fil.ion.ucl.ac.uk/spm). To correct for geometric distortions in the phase encoding direction, we calculated a deformation field based on reverse gradient estimation, using the Advanced Normalization Tools (ANTs) software package (http://stnava.github.io/ANTs/). In detail, we combined the mean functional volume (forward image) with the mean volume acquired with opposing phase encoding direction to generate a distortion-corrected template. Next, we estimated the deformation map by registering the forward image to the corrected template reference using non-linear (SyN) transformations. Finally, the deformation map was used to produce a distortion-corrected mean functional volume.

To co-register the anatomical and the distortion-corrected mean functional volumes, we initially referred to the Glasser’s atlas^113^ and the Kanwisher’s atlas^114^ to identify the approximate location of EVC and LOC, respectively. Then we outlined a volume containing these regions in the occipito-temporal cortex of both hemispheres on the individual participant’s native space (from here on referred to as manual mask) using ITK-SNAP with the 3D paintbrush tool (v.3.8)^115^. We then estimated a second deformation map in ANTs by registering the distortion-corrected mean volume to the T1-weighted volume, applying nonlinear (SyN) transformations within the manual mask. We visually inspected the fixed and registered volumes in ITK-SNAP for each participant. If the volume was not correctly registered within the region of the manual mask, we repeated the registration, adding linear transformations (rigid and affine) with stricter convergence criteria and increased iterations, until an accurate alignment was achieved. To minimize spatial resolution loss during resampling, we combined both deformation maps and resampled the functional images to the anatomical reference in a single interpolation step using a fifth-order spline function.

Finally, we spatially smoothed the functional localizer images using a 6-mm full width at half maximum (FWHM) Gaussian kernel. Functional images of the main runs were not smoothed to preserve spatial specificity.

Reliable layer analyses require accurate tissue segmentation. Because current automatic algorithms typically fail to capture fine cortical boundaries, we employed a manual segmentation approach to define the gray matter boundaries. Specifically, we delineated the outer and inner grey matter borders within the regions of interest (ROIs; see section “Definition of regions of interest” for details) on the T1-weighted UNI volume, following the procedure in ITK-SNAP outlined here (https://www.youtube.com/watch?v=tSA77mFTwcg&t=1042s). Prior to delineation, we corrected for bias field effects with a customized script^116^. The manually delineated gray matter surfaces were then used as input for automatic segmentation of the cortical ribbon into laminar and columnar compartments using the LN2_LAYERS algorithm from LAYNII (v2.2.1)^117^. In detail, we segmented the gray matter into three cortical depths within each ROI: deep, middle and superficial (**Fig. 3A**) applying the equi-distant model^39^. We chose the equi-distant model over the equi-volume model, as it provides more accurate layer segmentation for coarser resolutions (>0.3 mm), such as that of our fMRI data.

### fMRI univariate analysis

To estimate neural responses, we ran separate General Linear Model (GLM) analyses in SPM12 for each pre-processed functional run, i.e., all main experimental runs and the localizer run. All analyses were conducted in each participant’s native anatomical space. Specifically, we modelled 25 regressors (i.e., 24 object images + baseline) for each main experimental run, and 3 regressors (i.e., objects, scrambled objects, and baseline) for the localizer run. We created the regressors by convolving a boxcar function representing the onsets and durations of the corresponding condition with the canonical (2 Gamma) hemodynamic response function (HRF). We incorporated the motion estimates into the model as nuisance regressors. By fitting a GLM, we obtained beta weight estimates for each condition (i.e., regressor) for each run, which were subsequently used in further analyses.

### Definition of regions of interest

At the macroscale, we defined two ROIs for each participant: EVC, comprising areas V1, V2, and V3, and LOC in a two-step procedure. First, we used anatomical masks from the above-mentioned brain atlases for EVC^113^ and for LOC^114^. These masks were resampled from the MNI152 space into each participant’s individual space. Second, we identified the overlap between the participant-specific anatomical masks and the corresponding functional contrast T-statistic map from the localizer experiment, retaining the top 2,000 voxels. Specifically, we ranked the voxels according to the objects + scrambled > baseline contrast T-statistic for EVC, and the objects > scrambled contrast T-statistic for LOC. Due to lack of activation in the lateral occipitotemporal cortex for one participant (3), we used the objects + scrambled > baseline contrast to define LOC. Any voxels overlapping across ROIs were excluded. This process resulted in one final EVC and LOC mask for each participant.

At the mesoscale, we defined six ROIs based on the two brain regions – EVC and LOC – and the three cortical layers – deep, middle and superficial (see above for definition of cortical layers). Here, the ROI definition was analogous, with the following deviations to address issues arising specifically at the mesoscale level. Voxels with higher spatial resolution exhibit a reduced SNR and increased susceptibility to noise sources^47^, compromising the quality of the recorded signal. Additionally, signal reliability gradually decreases along the ventral visual cortex from lower to higher visual areas^118^, likely due to signal loss and susceptibility-induced distortions^119^. To address this, we chose a higher number of voxels for EVC compared to LOC. This resulted in 3000 and 1500 voxels with the highest T-statistic within the cortical ribbon of EVC and LOC, respectively. We then assigned each voxel to one of the three layers and selected those forming a complete column. For example, a voxel with column index 5 in the deep layer was only included if the corresponding voxels with column index 5 in the middle and superficial layers were also included.

### EEG data – paradigm, acquisition and analysis

We used a subset of the EEG data (*N* = 32) collected by Xie et al., 2024^5^ for the same images as in the fMRI experiment. Below is a summary of the relevant experimental procedures and acquisition steps.

The EEG experiment employed and backward masking paradigm consisting of 2,544 trials. In each trial, an object image was briefly presented for 17 ms and followed by a dynamic mask under two conditions: an early mask condition with inter-stimulus interval (ISI) of 17 ms or a late mask condition with an ISI of 600 ms. Object images and masks were randomly paired on each trial. All stimuli were centrally displayed on a gray background, with a size of 5° visual angle, and overlaid with a bull’s-eye fixation symbol. Participants were instructed to maintain fixation and refrain from blinking during trials, except during designated task trials (i.e., two-alternative forced-choice task, 25% of total trials) where blinking was allowed after making a response. The inter-trial interval (ITI) ranged from 900 – 1,100 ms; following the task trials, the ITI was extended to 2,000 ms to reduce motor artifacts.

For the present analyses, we focused on data from the late mask condition (ISI = 600 ms), yielding approximately 53 trials per object image. We analyzed the time window from 200 ms pre-stimulus to 600 ms post-stimulus, before mask onset.

EEG data were recorded with a 64-electrode ActiCap system and a Brainvision actiChamp amplifier. Electrodes were positioned according to the international 10-10 system, with a ground electrode and a reference electrode placed on the scalp. The signals were sampled at 1,000 Hz and online filtered between 0.03 and 100 Hz. EEG data were preprocessed using Brainstorm-3^120^. Noisy channels (mean = 2.2, SD = 1.8) were removed, and the data were low-pass filtered at 40 Hz. Independent component analysis was applied to remove eye movement and other artifact components (mean = 2.7, SD = 0.9). Data were then segmented into epochs from −200 ms to 600 ms relative to stimulus onset, baseline-corrected, and multivariate noise normalization^121^ was applied for decoding analyses.

Multivariate analysis was performed on a participant-specific basis using SVMs as implemented in the LIBSVM toolbox in MATLAB (2021a). To determine when the brain processes object information, time-resolved decoding analysis was conducted from −200 ms to 600 ms relative to target image onset in 10 ms intervals. At each time point, trial-specific EEG channel activations were extracted and arranged into 64-dimensional pattern vectors for each of the 24 object image conditions. For each condition, trials were randomly grouped into four equally sized bins and averaged to create four pseudo-trials, repeated over 100 permutations. Pseudo-trials were then divided into a training set (three pseudo-trials) and a testing set (one pseudo-trial) for pairwise object identity decoding. This process was repeated for all pairwise combinations of object conditions.

### Multivariate pattern analysis of fMRI data

To decode object information from voxel activation patterns in EVC and LOC, we used SVMs as implemented in the scikit-learn library in Python. Analyses were conducted separately for each participant and for EVC and LOC at the macroscale. We aggregated run-specific voxel-wise beta estimates derived from the GLM into pattern vectors for each of the 24 conditions. To enhance the signal to noise ratio, we averaged beta weights across two randomly selected subsets of trials into two pseudo-trials for each condition. We repeated this process 300 times, each time performing pairwise object decoding, for all object condition combinations, using a two-fold cross-validation approach. We averaged the results across all iterations and all object condition combinations to yield one grand-average decoding accuracy for each ROI and participant.

### Representational similarity analysis

To relate object representations across different signal spaces (i.e., voxel activation patterns in fMRI, sensor activation patterns in EEG, embeddings in DNNs), we used representational similarity analysis^25^. This approach is based on the rationale that if two images elicit similar neural representations, their corresponding fMRI, EEG or DNN signal patterns should also be similar. We used two variants of RSA: (i) representational similarity analysis-based fusion^25,27,28^, relating spatially localized object representations (from fMRI) to specific temporal dynamics (from EEG) to resolve the spatiotemporal dynamics with which visual representations emerge, and (ii) representational similarity analysis-based commonality analysis^122,123^ to determine the visual feature complexity (from layer-specific DNN embeddings) of spatiotemporally identified dynamics. For this we calculated the common variance between spatially localized object representations (from fMRI), specific temporal dynamics (from EEG), and layer embeddings (from DNNs). We detail both approaches below after describing the specifics of summarizing representational similarity for each signal space in representational dissimilarity matrices (RDMs).

### Construction of fMRI representational dissimilarity matrices

To construct the fMRI-RDMs, we first extracted and vectorized beta weight estimates for each of the 24 conditions to form fMRI neural patterns for a given ROI (area or cortical layer). To account for voxel-wise differences in noise and signal variance across layers, we applied multivariate noise normalization^124,125^. To quantify pattern relationship while mitigating scaling effects introduced by draining veins^40^, we used Pearson’s correlation as a scale-invariant measure for all pairwise combinations of experimental conditions. Output correlations were transformed into dissimilarity values using 1 – Pearson’s correlation, and organized into a 24×24 fMRI representational dissimilarity matrix (RDM), indexed in rows and columns by the compared conditions. For each participant, this yielded two fMRI-RDMs at the macroscale level of brain areas: the EVC RDM and LOC RDM; and six layer-specific fMRI-RDMs at the mesoscale level of cortical layers: Deep EVC RDM, Middle EVC RDM, Superficial EVC RDM, and Deep LOC RDM, Middle LOC RDM and Superficial LOC RDM. To contextualize the model performance with respect to noise in the fMRI measurement, we computed the noise ceiling (**Suppl. Fig. 4D, E and F**). Given the higher noise levels in fMRI, particularly in layer-resolved measurements, compared to EEG, we followed previous EEG-fMRI fusion studies^5,26,30^ and averaged the fMRI-RDMs across participants to optimize signal quality.

### Construction of EEG representational dissimilarity matrices

A detailed description of the creation of the EEG-RDMs is provided by Xie et. al., (2025)^5^. Briefly, for each participant and each time point in the epoch from −200 to 600 ms, we pairwise decoded object information using EEG channel activation patterns, via SVMs as implemented in the LIBSVM toolbox^126^ in MATLAB (2021a). The decoding accuracies for each pair of object images were assembled into 24×24 EEG-RDM for each time point (801 in total), with rows and columns indexed by the conditions. Each matrix was symmetric across the diagonal, with diagonal entries left undefined.

### Construction of DNN representational dissimilarity matrices

To compute the DNN-RDMs, we used the Vision Transformer architecture^35^ from the DeiT family^36^, specifically the deit_small_patch16_224 model trained on the ImageNet dataset^127^ for visual object categorization. In detail, we presented the same set of 24 object images to the pretrained network and extracted the activation patterns for each object condition from the final output embeddings of the first, seventh, and twelfth transformer encoder blocks, as well as from the final classification head. Next, we quantified the pairwise pattern dissimilarity using 1 – Pearson’s correlation for all pairs of image combinations and organized the outputs into 24×24 DNN-RDMs, with rows and columns representing Pearson’s based dissimilarity measure between object conditions. This resulted in a total of 7 layer-specific DNN RDMs.

### Representational similarity analysis-based EEG-fMRI fusion

We used representational similarity analysis-based fusion^25,27,28^, relating fMRI-RDMs to EEG-RDMs using Spearman rank order correlation. We applied this analysis at the macroscale and at the mesoscale, for each participant separately. At the macroscale, we related region-specific RDMs (i.e., EEG, LOC) to EEG-RDMs, yielding one time course for each ROI for each participant. At the mesoscale, to control for non-specific macrovascular responses^40,41^, we performed EEG-fMRI fusion analyses in which, for each cortical layer, we partialed out the influence of the underlying layers using partial Spearman rank order correlation. Specifically, for the superficial layer, we partialed out the effect of the middle layer. Similarly, for the middle layer, we partialed out the effect of the deep layer. The deep layer remained unaffected by this approach. By relating region- and layers-specific ROIs to EEG-RDMs, we obtained one time course for each of the six ROI-layer combinations for each participant.

### Representational similarity analysis-based commonality analysis

To characterize the level of feature complexity of the neural representations identified above in space and time, we related them to DNN embeddings across model layers. Feature complexity increases progressively across DNN layers: early model layers are more sensitive to low complexity features, whereas later layers are tuned to high-complexity features^128,129^, analogous to the ventral visual hierarchy in humans and non-human primates^34,38^. Thus, by linking the neural representations to those from the DNN layers, we can determine the feature complexity of spatially localized object representations to particular time points. We did this using representational similarity analysis-based commonality analysis^110^. In detail, we calculated the commonality coefficient corresponding to the shared variance among fMRI-RDMs for each brain region and cortical layer, EEG-RDMs at time points with significant effects in EEG-fMRI fusion (averaged across time), and DNN RDMs for each model layer.

### Quantification and statistical analysis

To assess statistical significance, we performed non-parametric statistical analyses that do not rely on assumptions about the data distribution. The empirical estimations, including decoding accuracy as well as correlation values from the representational similarity analysis-based fusion and from the commonality analysis, were tested against a null distribution created by sign permutation (1,000 permutations). We randomly multiplied the participant-specific data by ±1 to generate permutation samples, recomputed the statistic for each sample, and derived *P* values. All permutation tests were one-sided (right-tailed), consistent with the directional hypothesis that the observed effects were greater than zero.

We controlled the familywise error rate across time points using cluster-based statistics. First, *P*-value maps were thresholded at *P* < 0.05 to define temporally contiguous suprathreshold clusters. These clusters were then used to construct an empirical null distribution of maximum cluster weights, and a corrected threshold was determined at the 95th percentile of the right tail of this distribution. For the commonality analyses we corrected *P* values for multiple comparisons by FDR-correction.

We estimated the 95% confidence intervals for the peak latencies in the RSA-derived time courses using bootstrapping (1,000 paired bootstrap resamples with replacement). For each bootstrap we calculated the statistic, resulting in a bootstrap estimate of peak latencies from which we derived the confidence intervals.

To estimate confidence intervals for peak latency differences, we used an analogous bootstrapping approach, resampling the mean peak-to-peak latency difference for each resample. This generated a distribution of mean differences, from which we derived the 95% confidence interval. We set *P <* 0.05, i.e., if the 95% confidence interval did not include 0, we rejected the null hypothesis of no peak-to-peak latency differences.

## Supplementary information

**Supplementary Figure 1.**
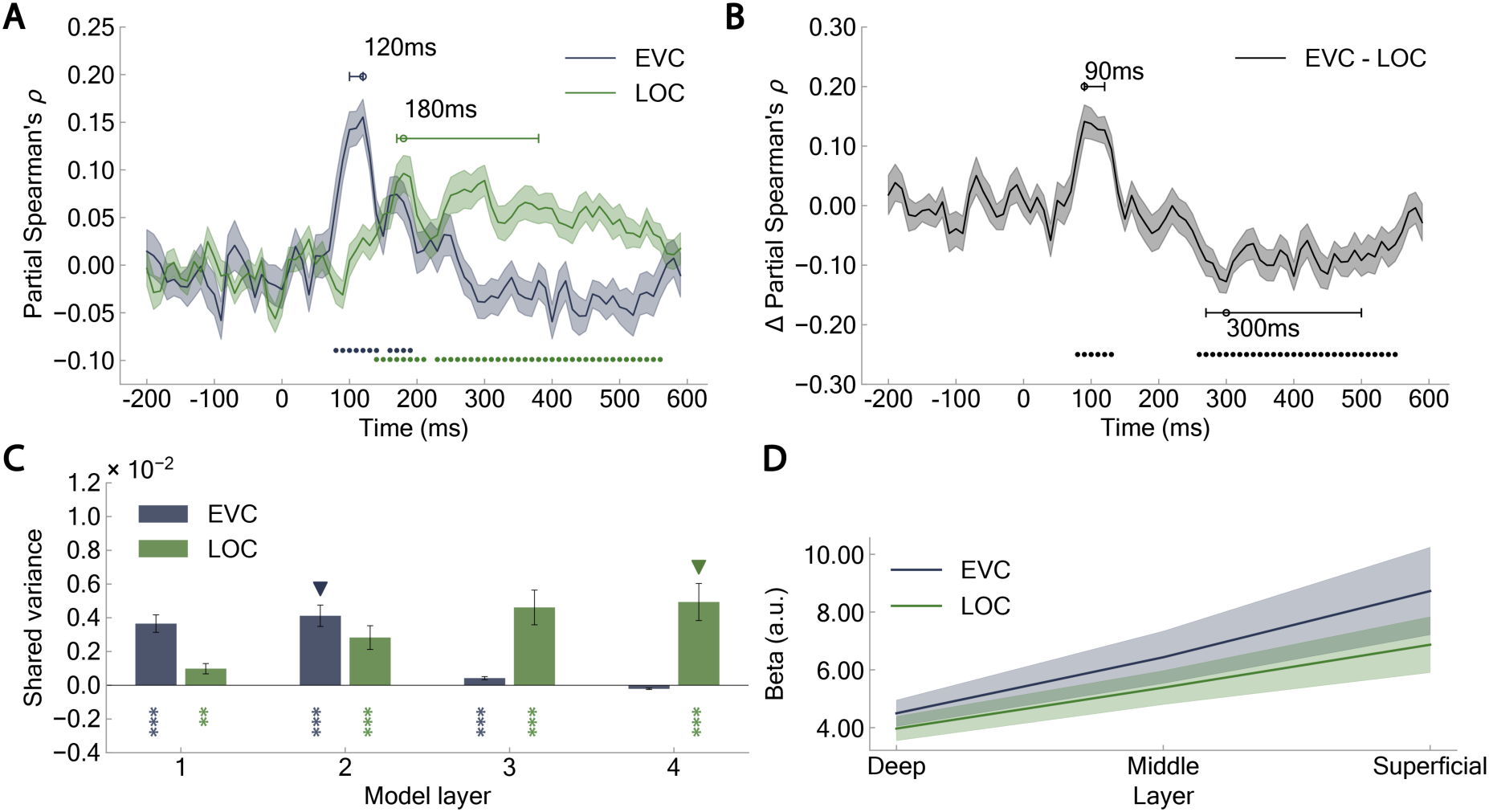
Macroscale representational EEG-fMRI fusion and commonality analysis using partial correlations, and laminar profiles of the GE-BOLD response. (**A**) Spatiotemporal neural dynamics at the macroscale level. For EVC we partialed out the effect of LOC and for LOC we partialed out the effect of EVC. Early representations correlated more strongly with EVC than with LOC and later representations correlated more strongly with LOC than with EVC. (**B**) Difference between EVC and LOC curves in (**A**). (**C**) Macroscale commonality analysis for time intervals showing significant EVC–LOC differences. Visual representations of low-to-mid-complexity emerge primarily in EVC, while mid-to-high-level object representations emerge in LOC. (**D**) Laminar profiles of the GE-BOLD response to objects in EVC and LOC. Laminar responses derived from GE-BOLD signals are strongly affected by non-specific macrovascular signals, leading to higher activation in superficial layers. Data were averaged across 24 object conditions and 10 fMRI participants. Shaded area represents the standard error of the mean across EEG participants. Shaded regions denote the standard error of the mean across participants; colored circles indicate significant time points (*N* = 32, cluster-defining threshold *P <* 0.05, cluster threshold *P <* 0.05); uncolored circles and horizontal lines indicate peak latency means and 95% confidence intervals, respectively. Colored asterisks indicate significant correlations (*N* = 32, right-tailed permutation tests, FDR-corrected; \**P <* 0.05; \*\**P <* 0.01; \*\*\**P <* 0.001); colored triangles represent model layers with the highest occurrence proportion, determined through 1,000-iteration bootstraps.

**Supplementary Figure 2.**
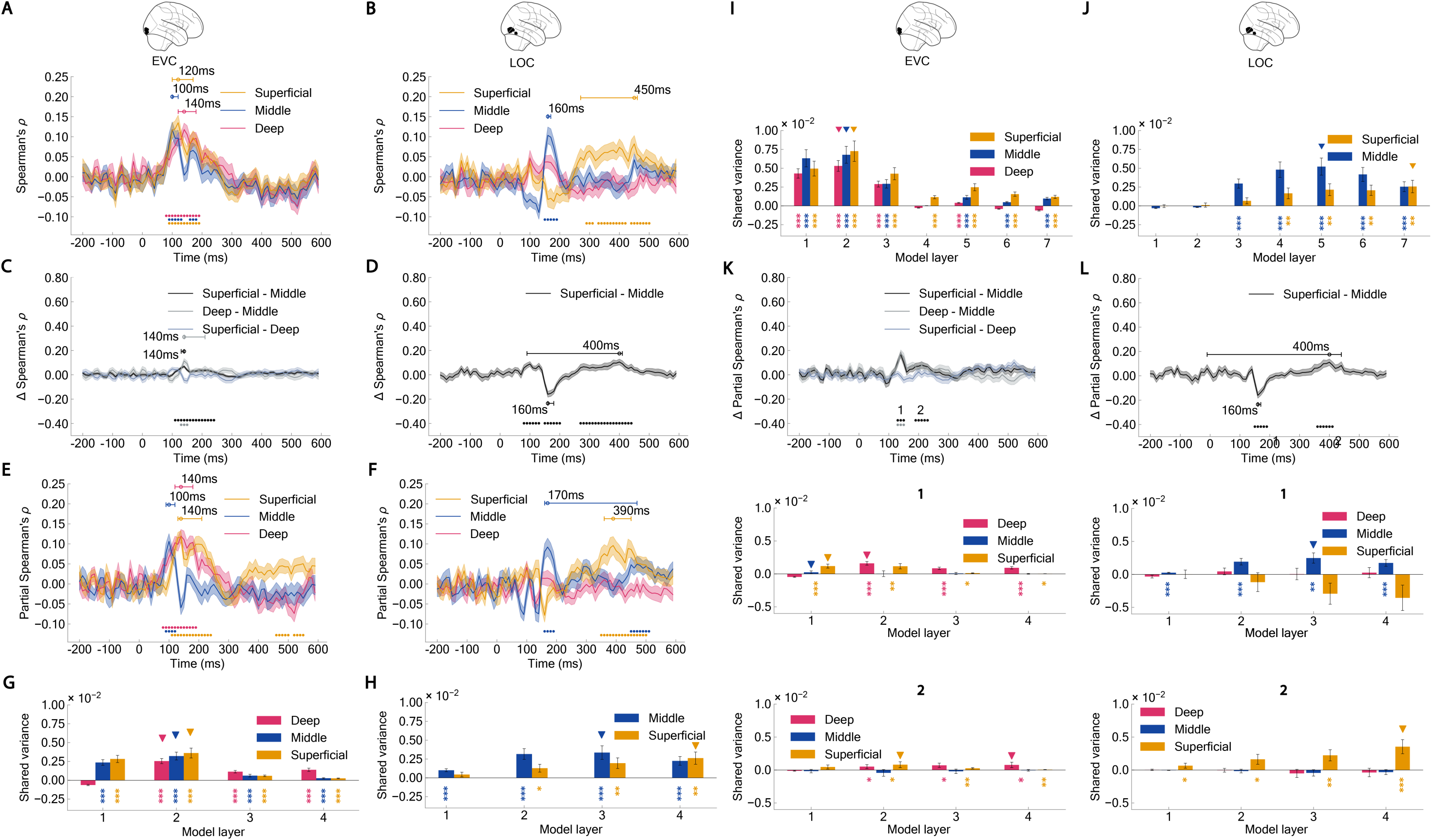
Mesoscale representational EEG–fMRI fusion and commonality analyses under alternative experimental choices. (**A**–**D**) Representational EEG–fMRI fusion without partial correlations. (**A**) Layer-specific spatiotemporal neural dynamics in early visual cortex (EVC). (**B**) Differences between EVC layer-specific curves shown in (**A**). (**C**) Layer-specific spatiotemporal neural dynamics in LOC. (**D**) Differences between LOC layer-specific curves shown in (**C**). (**E**–**H**) Representational EEG–fMRI fusion and commonality analyses with matched voxel counts across cortical layers. For each layer, voxels were randomly subsampled to match the layer with the fewest voxels; this procedure was repeated ten times and results were averaged across permutations. (**E**) Layer-specific spatiotemporal neural dynamics in EVC. (**F**) Layer-specific spatiotemporal neural dynamics in LOC. (**G**,**H**) Representational format (≈ feature complexity) across cortical layers in EVC and LOC, respectively. (**I**,**J**) Representational format (≈ feature complexity) across cortical layers in EVC and LOC using AlexNet, respectively. (**K**,**L**) Representational format (≈ feature complexity) across cortical layers in EVC and LOC for time intervals showing significant between-layer differences. Two temporal clusters (labeled 1 and 2) were defined from these intervals. For each cluster, commonality analysis linked model layer-specific DNN-RDMs to layer-specific fMRI-RDMs and the mean EEG-RDM within the corresponding time window in EVC (**K**) and LOC (**L**). Shaded regions and error bars indicate the standard error of the mean across participants. Colored circles denote significant time points (*N* = 32; cluster-defining threshold *P* < 0.05; cluster threshold *P* < 0.05); uncolored circles and horizontal lines indicate peak latency means and 95% confidence intervals, respectively. Colored asterisks indicate significant correlations (right-tailed permutation tests, FDR-corrected; \**P* < 0.05; \*\**P* < 0.01; \*\*\**P* < 0.001). Colored triangles represent model layers with the highest occurrence proportion, determined through 1,000-iteration bootstraps.

**Supplementary Figure 3.**
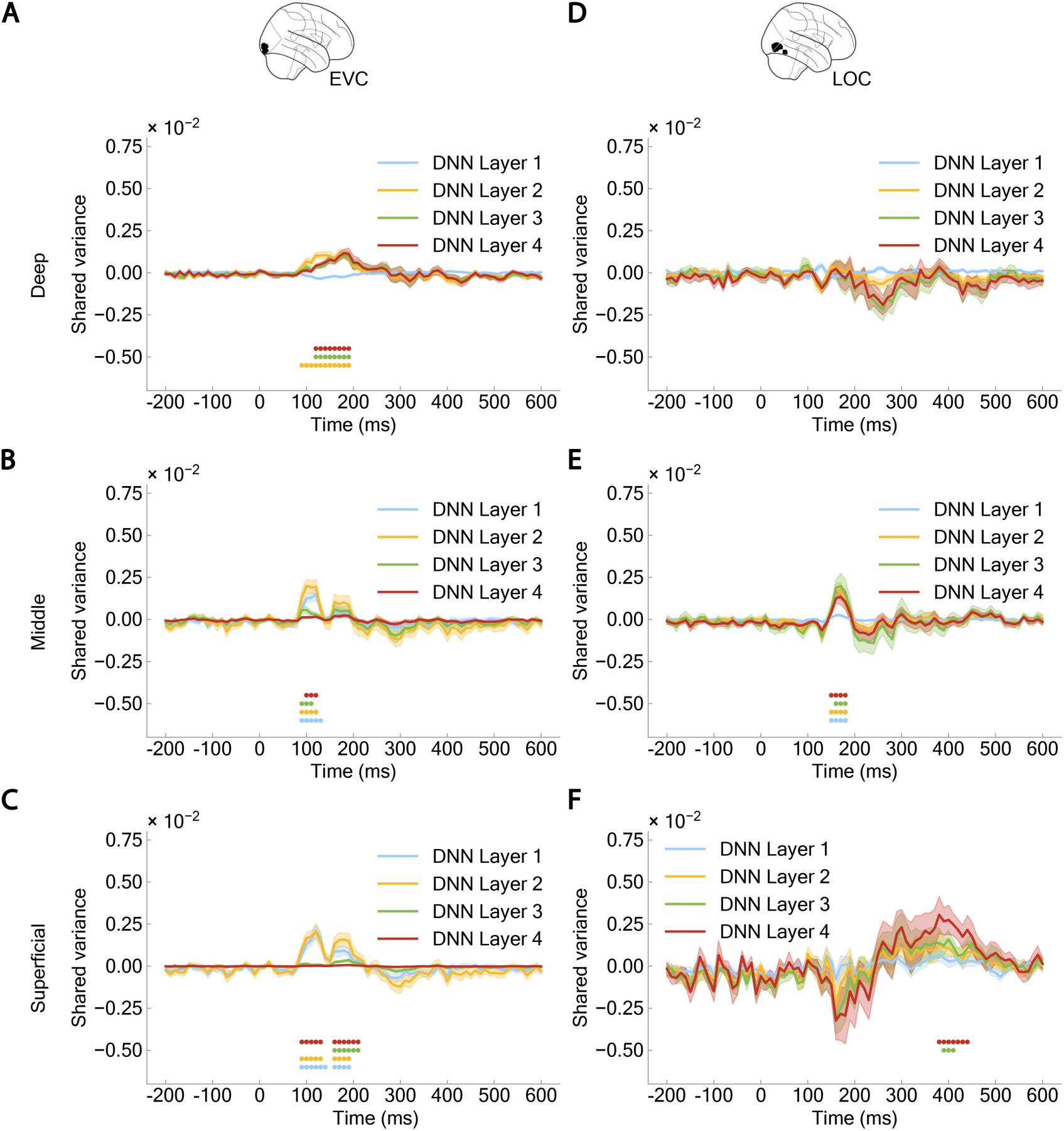
Mesoscale commonality analysis across individual time points. Representational format (≈ feature complexity) in deep layer (**A**, **D**), middle (**B**, **E**), and superficial (**C**, **F**) layers in EVC (**A–C**) and LOC (**D–F**). Shaded regions denote the standard error of the mean across participants. Colored circles indicate significant time points (*N* = 32, cluster-defining threshold *P <* 0.05, cluster threshold *P <* 0.05).

**Supplementary Figure 4.**
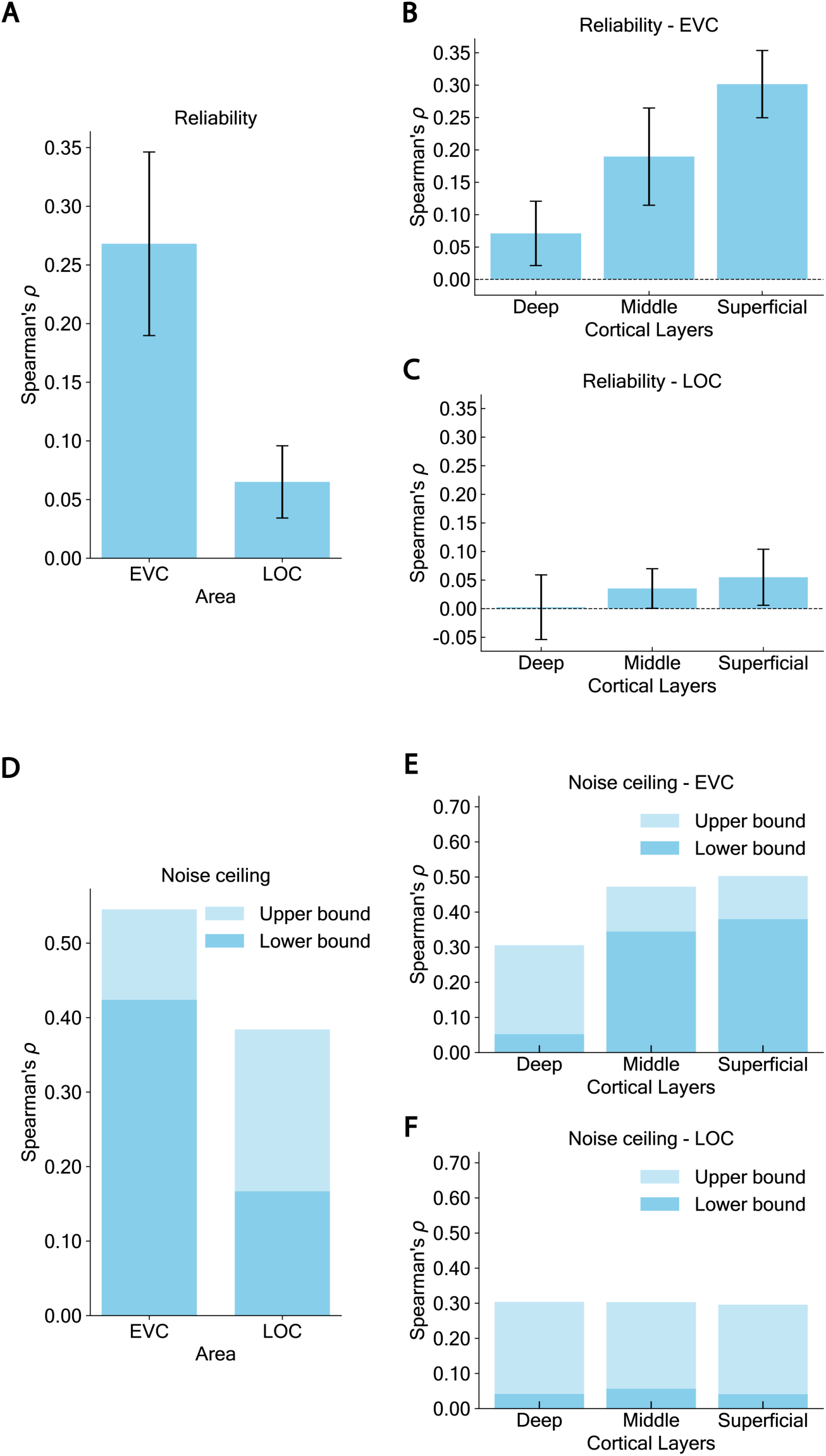
Reliability and noise ceiling of the fMRI data at macroscale and mesoscale. RDM reliability at the macroscale across brain regions (**A**) and at the mesoscale across cortical layers in EVC (**B**) and LOC (**C**). Noise ceiling estimates at the macroscale (**D**) and mesoscale in EVC (**E**) and LOC (**F**). Error bars indicate the standard error of the mean across participants (*N* = 10).

